# An atlas of positive selection in the genomes of major malaria vectors

**DOI:** 10.1101/2025.07.16.664900

**Authors:** Sanjay C Nagi, Nicholas J Harding, Mara K. N. Lawniczak, Martin J Donnelly, Alistair Miles

## Abstract

Insecticide resistance poses a major threat to malaria control efforts in Africa, yet our understanding of the genomic basis of adaptation in malaria vectors remains incomplete. Here we present a comprehensive atlas of recent positive selection in natural populations of the major African malaria vectors; the *An. gambiae* species complex and *An. funestus*. By analyzing whole-genome sequence data from 4,306 mosquitoes collected across twenty-one countries, we identify and characterize both known and novel genomic regions under strong positive selection. We develop an innovative approach for identifying and localising selection signals, and find extensive evidence for parallel evolution between species, with all important loci in *An. funestus* also under selection in *An. gambiae s.l.* We confirm intense selection at established resistance loci including known target site proteins (*Vgsc, Rdl, Ace1*) and metabolic enzymes (*Cyp6p*, *Cyp9k1*, *Gste*, *Coeaexf*) and describe novel signals at a diacylglycerol kinase on the X chromosome, and in *An. gambiae s.l*, at the metabolic regulator *Keap1*. To support real-time monitoring of emerging variants threatening vector control, we present an open-source web resource containing detailed selection scan results for all cohorts. The resource is driven by an automated computational workflow, which enables continuous updates as new whole-genome data are generated. The selection atlas provides a foundation for understanding vector adaptation to insecticide-based interventions and we hope will inform evidence-based resistance management strategies that are critical for sustaining malaria control.

## Introduction

The environment in which African malaria mosquitoes live and breed has been radically altered in recent times. In the last 30 years the total human population of sub-Saharan Africa has more than doubled, with urban populations tripling in size^1^. This rapid growth has expanded niches for anthropophilic mosquitoes that are able to adapt to the built environment^2,3^. Human population growth has created a demand for increased agricultural productivity, which has been achieved largely by cultivating more land, supported by the introduction and widespread use of chemical pesticides^4^. In the last two decades, massive public health campaigns have been mounted to reduce the burden of malaria, via mass distribution of long-lasting insecticidal bed nets (LLINs) and large-scale programmes of indoor-residual spraying (IRS) of insecticides^5^. In the face of these extraordinary selective pressures, mosquito populations are known to be evolving rapidly, and resistance to public health insecticides has become almost ubiquitous^6^.

Many of the adaptations involved in insecticide resistance in Anopheles mosquitoes are well characterised. The Voltage-gated sodium channel (*Vgsc)* is the physiological target of both DDT and pyrethroid insecticides, and substitutions within the protein have been shown to interfere with insecticide binding, conferring target-site resistance^7,8^. In addition, a combination of amino acid mutations and copy number variation has been found in *Ace1*, a gene encoding acetylcholinesterase, the target of carbamate and organophosphate insecticides^9^. The *Rdl* locus encodes a subunit of a GABA-gated chloride channel, the target of dieldrin, an organochlorine insecticide used heavily in both agriculture and public health during the 1950s and 1960s. Two variants in *Rdl* that cause resistance to dieldrin have been found in *Anopheles* populations and persist despite a ban on dieldrin in the 1970s^10,11^.

Detoxifying enzymes have also been implicated in adaptation to insecticides across a range of insect taxa^12^. Three major protein families are primarily involved in metabolic detoxification: cytochrome P450s, glutathione S-transferases, and carboxyl or alpha-esterases^13^. Genes in these families are commonly found to be over-expressed in resistant mosquitoes^14^, commonly driven by extensive copy number variation^15,16^ or other regulatory mutations, such as transposable element insertions^17,18^. Amino acid mutations in metabolic enzymes that increase the efficiency of insecticide metabolism have also been reported, for example, a point substitution in the *Gste2* gene enhances the metabolism of DDT in both *An. gambiae s.l* and *An. funestus*^19,20^.

Several studies have reported apparent changes in mosquito behaviour, whereby mosquitoes avoid contact with insecticides inside homes by feeding outdoors or at times when individuals are not protected by LLINs^21^. Evidence for adaptation rather than behavioural plasticity remains equivocal, and no genetic associations have yet been found with behavioural differences, although several olfactory genes were reported as commonly over-expressed in a recent meta-analysis^14^. Adaptations that alter the mosquito cuticle such that insecticides are less able to penetrate have also been reported in multiple *Anopheles* species^22–25^. Additionally, there is evidence of involvement in resistance of some genes from less characterised gene families, such as chemosensory proteins^14,26,27^.

In response to the widespread level of resistance driven in part by two decades of pyrethroid-only LLIN use, the vector control armamentarium has changed in recent years, with next-generation LLINs coming to market. In 2024, dual active-ingredient nets comprised 51% and Piperonyl butoxide (PBO) - pyrethroid nets 33% of all LLINs delivered in sub-Saharan Africa^28^. These novel tools have shown superior epidemiological efficacy in randomized controlled trials^29,30^, however their widespread deployment raises concerns about repeating earlier mistakes of over-reliance on single classes of chemistry. To ensure the longevity of these products, the vector control community must employ them strategically as part of coordinated insecticide resistance management practices. These may include mosaics that alternate different active ingredients between areas, temporal rotation of different interventions, and mixture strategies combining multiple modes of action^31^. Additionally, integrating and advancing non-chemical methods such as house screening and larval source management could help to both reduce the malaria burden and the selection pressure from insecticides.

Genomic surveillance will be crucial for detecting and tracking novel resistance variants before they become widespread^32^. While whole-genome sequencing remains vital for discovering new resistance loci through genome-wide association studies, targeted amplicon sequencing enables cost-effective routine monitoring at known resistance markers. Recent developments in laboratory protocols and computational pipelines now provide standardized frameworks for real-time genomic surveillance^33^. These tools allow rapid detection of resistance mutations and enable evidence-based decision making about intervention deployment.

Here we analyze whole-genome sequence data from 4,306 mosquitoes collected across twenty-one African countries as part of the *Anopheles gambiae* 1000 Genomes Project (Ag1000G), the *An. funestus* 1000 genomes project, and the Vector Observatory^34,35^. The data comprise two separate cohorts of high-quality genotypes and haplotypes from the *An. gambiae* species complex and *An. funestus s.s*. We present these results through an interactive web resource - the malaria vector selection atlas - providing comprehensive selection scan results and highlighting emerging threats to vector control. The resource is powered by an automated Snakemake workflow enabling continuous updates as new data are available.

## Results

### Sample manifest

The dataset contains high-coverage (30x) whole-genome sequence data from 4306 individual *Anopheles gambiae s.l* (3763) and *An. funestus* (543) mosquitoes collected in twenty-one countries across sub-Saharan Africa. Figure 1 shows the sampling locations of each cohort. Individual mosquitoes were collected as adults or larvae, between 1994 and 2018 (see Supplementary Table 1). We aggregated mosquitoes into cohorts based on the geographic district and the quarter of the year in which they were collected. We included a cohort in our analysis if there were ≥15 individuals. For computational efficiency, if there were more than 200 mosquitoes in a given cohort, we randomly downsampled their cohort to 200. This resulted in 63 cohorts from the *An. gambiae* complex and 15 from *An. funestus*.

**Figure 1.**
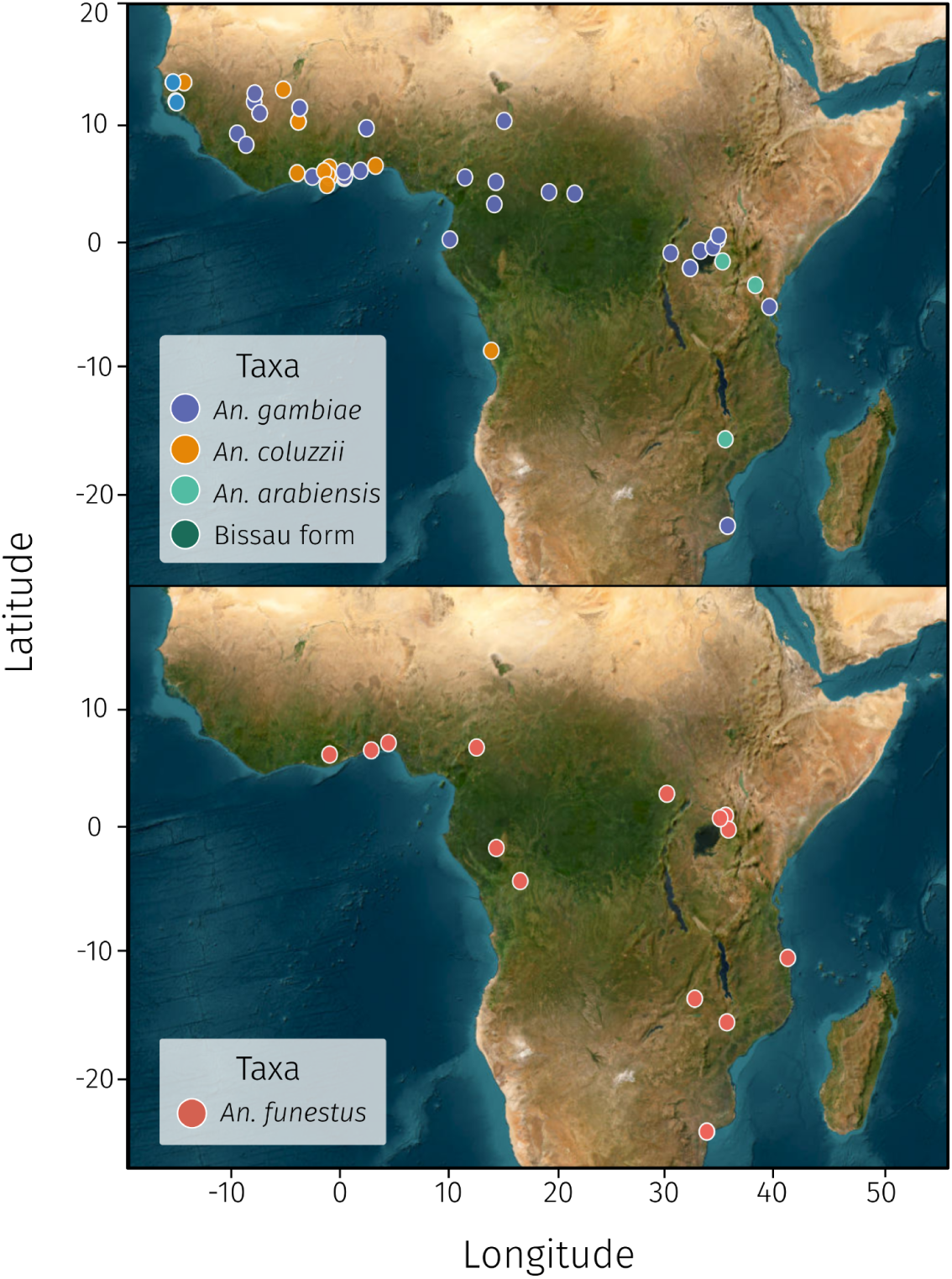
Geographic distribution of Anopheles mosquito samples. Maps showing the sampling locations of cohorts of *An. gambiae s.l.* (upper) and *An. funestus* (lower) mosquitoes collected across 21 countries in sub-Saharan Africa as part of the *An. gambiae* 1000 Genomes Project phase 3 and the *An. funestus* 1000 genomes project phase 1. Samples were collected as adults or larvae between 1994 and 2018 and grouped into cohorts based on geographic district and collection period.

### Genome-wide selection scans

For each cohort in our data, we computed three genome-wide selection statistics, H12, G123 and iHS. H12 provides a measure of haplotype diversity - a high value of H12 indicates an unusually low haplotype diversity, expected in genomic windows where recent selective sweeps containing a beneficial allele have spread to a moderate or high frequency^36^. G123 is analogous to H12 but applied to unphased diplotypes rather than haplotypes^37^, included for situations where phasing may be challenging. We also computed the integrated haplotype score (iHS) for each cohort, a statistic that identifies incomplete selective sweeps by comparing the lengths of haplotypes associated with alternative alleles at a given SNP, with longer haplotypes indicating recent positive selection.

### Signal calling

Selection signals observed at known insecticide resistance genes not only showed extreme outlying values across the three statistics, but also had a clear peak architecture, with values of each decaying on both flanks. This is a well-known phenomenon in selection analyses due to genetic hitchhiking of neighbouring loci^38,39^. To call signals within an automated workflow, we developed an approach to exploit this signal decay. Several previous studies have used simple heuristic peak-finding algorithms for locus discovery^36,40,41^, however, these approaches may result in a lack of sensitivity where true signals are close to one another. We developed an algorithm based upon the application of exponential decay curves to the data using non-linear least squares regression^38^ to systematically search the H12 data in windows for peak-like signals. We fitted an exponential decay curve, a Gaussian curve, and a null constant straight-line model to windows across the genome. In each case, we evaluated the difference in model fit (Δi) between the best-fitting alternate model and a null constant model. We recorded the window with the highest Δi, then subtracted the fitted values from the dataset, allowing the discovery of subsequent peaks in an iterative process, until we reached our confidence threshold (> 1000 Δi). Supplementary Data 2 contains a full table of the signals detected.

### Web resource

To make our findings accessible to the vector biology and control community, we developed an interactive web-based resource. The selection atlas (https://anopheles-genomic-surveillance.github.io/selection-atlas/) presents the results of our genome-wide selection scans across all sampled cohorts and features interactive visualizations that allow users to explore selection signals by chromosome, country, or individual cohort. The resource highlights both known and novel genomic regions under putative selection through a curated series of “selection alerts” that identify loci of public health concern. Each alert page provides detailed information about the genomic region, affected cohorts, and candidate resistance genes with supporting evidence.

### Selection alerts

#### Selection signals at known resistance loci

To obtain an empirical sense of expected statistic values at previously validated signals of selection, we examined the results of our selection scans at genome regions containing genes known to play a functional role in insecticide resistance.

#### Vgsc

The Voltage-gated sodium channel gene (*Vgsc*) is the physiological target of both DDT and pyrethroid insecticides, and amino acid substitutions in codon 995 from Leucine to Phenylalanine^7,42^ or Serine^43^ are known to cause knock-down resistance (*Kdr*) via conformational changes in the sodium channel^44^. Phenylalanine mutations in homologous codons have been described in numerous pest species under selective pressure from DDT and pyrethroids^8,45^. In *An. gambiae*, haplotype network analyses have revealed at least five distinct origins of the L995F allele (F1-F5) and five for L995S (S1-S5), with evidence that several haplotype groups have spread across thousands of kilometres^46^. Both codon 995 alleles show widespread distribution across Africa, with evidence of introgression of a 995F haplotype from *An. gambiae* into *An. coluzzii*^47^. Secondary non-synonymous mutations have arisen on L995F haplotypes, including N1570Y that significantly enhances resistance, and others such as P1874S/L and T791M, which may enhance resistance or compensate for fitness costs. In contrast, few secondary mutations have been observed on L995S backgrounds. A novel combination of mutations has been identified in West African *An. coluzzii*, comprising I1527T in tight linkage with V402L^46^. This is particularly significant as I1527T occurs adjacent to the predicted pyrethroid-binding site, and substitutions orthologous to V402L have been shown to confer resistance in other insect species^48^. In agreement with the earlier literature, the *Vgsc* locus shows pronounced selection in 37 cohorts across West Africa (Benin, Ghana, Guinea, Mali, Côte d’Ivoire), Central Africa (Cameroon, DR Congo), and East Africa (Uganda, Tanzania). Both *An. gambiae* and *An. coluzzii* exhibit these signals, with H12 values reaching 1 in many populations, meaning that only one or two haplotypes are now segregating this locus. No signals were found at this locus in *An. arabiensis* in these data. *Vgsc* occurs on chromosome arm 2L at approximately position 2.4 Mbp in a pericentromeric region, where a lower recombination rate is likely to have caused some asymmetrical distortion in this signal.

Knockdown resistance at the *Vgsc* in *An. funestus* is not widespread and had not been detected until a recent study from Tanzania, found the *Vgsc*-995F mutation at high frequencies in Morogoro district with an accompanying selection signal, but absent from other areas of the country^49^. This discovery was then linked to historic stockpiles of DDT that had contaminated ground soil. In this dataset, however, no signals were observed at the locus - the Morogoro cohort was excluded due to having only 10 individuals in total.

#### Ace1

Carbamate and organophosphate insecticides inhibit the action of acetylcholinesterase, causing insect death by blocking synaptic neurotransmission. Acetylcholinesterase is encoded by the *Ace1* gene (AGAP001356) in *An. gambiae*, variation in which has been identified as conferring resistance. The G280S mutation (G119S in *Torpedo californica)* results in an altered protein structure, affecting binding affinity of the insecticide^50,51^. This mutation is common in *An. gambiae* throughout Western Africa^9^. In a highly resistant *An. coluzzii* population from Tiassale, Côte d’Ivoire, *Ace-1* is significantly upregulated, mediated at least partially by a duplication at this locus ^52^. Selection at the acetylcholinesterase locus appears more geographically confined than other loci, with signals in just 5 cohorts all from West Africa (Ghana, Côte d’Ivoire). The signals predominantly occur in *An. gambiae* with some representation in *An. coluzzii*. H12 values peak at 0.56 in these cohorts. Although clearly defined peaks were visible around this gene, scan values (Δi = 1269-2920) were less extreme than at other known loci, which could be due to gene duplications affecting our ability to phase haplotypes accurately. Indeed, the H12 data exhibits a slight dip in the middle of the signal, in contrast to G123, which is applied to unphased diplotypes. In *An. funestus*, the *Ace1-*G119S is absent, however, another mutation, *Ace1*-N485I, has been detected at low frequencies in southern Africa, and has been shown to confer resistance to carbamates^53^. No signals were detected in these data, nor is N485I.

#### Rdl

Variation in the γ-aminobutyric acid (GABA) receptor in the Rdl gene causes target site resistance to the insecticide dieldrin. Dieldrin resistance has been attributed to single amino acid substitutions in *D. melanogaster*^54^ and other pest species^55,56^. Two resistance mutations have been found in anophelines, both in codon 296: alanine-to-glycine (A296G) and alanine-to-serine (A296S)^10^. The A296G allele is common in West and Central African populations of both *An. gambiae* and *An. coluzzii*, while A296S is found in Central and East Africa. These resistance alleles are associated with nearby mutations that likely provide compensatory effects, particularly T345M with A296G and T345S with A296S. Despite barriers to gene flow, these resistance alleles spread through rare combinations of interspecific introgression (from *An. gambiae* and *An. arabiensis* to *An. coluzzii*) and across different karyotypes of the 2La chromosomal inversion. The 296G allele underwent introgression from *An. gambiae* to *An. coluzzii* and later spread from 2L+a to 2La chromosomes^10^. The 296S allele originated in *An. arabiensis* and later introgressed into *An. coluzzii*. *An. gambiae* and *An. coluzzii* show selection signals spanning West Africa (Benin, Burkina Faso, Ghana, Mali, The Gambia) and Central Africa (Cameroon, Central African Republic), across 23 cohorts overall. Remarkably, a cohort from Ghana (GH-AA_La-Nkwantanang-Madina_gamb_2017_Q4) exhibits H12 values of 0.97, which may be suggestive of continued selection pressure on this population since the ban of dieldrin around 50 years ago.

Similarly to *An. gambiae*, a A296S mutation is found in the GABA receptor in *An. funestus*^11,35^. Selection at this locus is also widespread, observed in populations from Ghana, Nigeria, and Uganda (peak H12 = 0.7). Dieldrin was banned in the 1970s due to its high persistence as an organic pollutant and unexpectedly wide toxicity. Despite this ban, resistance alleles have remained strikingly persistent in natural *Anopheles* populations for over 40 years.

#### Cyp6 cluster

Cytochrome P450 enzymes play a range of roles in insects and are important mediators of insecticide resistance in *An. gambiae*, being the major gene family responsible for the metabolism of pyrethroids. Of the 111 P450 genes in *An. gambiae*, only members of the subfamily CYP6 and CYP9 have been shown to be capable of metabolising pyrethroids. *Cyp6aa1*, *Cyp6p3* and *Cyp6m2* among others have been frequently identified in gene expression studies as being over-expressed in resistant mosquitoes^18,57,58^. Additionally, upregulation of P450s confers cross-resistance to other insecticides in *An. gambiae*^52^ and other insect species^59^. A cluster of the Cyp6 subfamily is located on chromosome 2R, around 28.5 Mbp, containing several *Cyp6p* genes. This enzyme cluster exhibits the most extensive footprint selection in the dataset, with 48 cohorts showing signals across the entire sampling range. *An. gambiae, An. coluzzii, An. arabiensis* and a cryptic taxon, the Bissau molecular form, display strong sweeps with extremely high H12 values common across cohorts and some of the highest Δi values in the dataset (up to 8618).

The *Cyp6p* cluster exhibits particularly strong selection signals in *An. funestus* populations across sub-Saharan Africa, with cohorts from nine countries (Benin, Cameroon, Democratic Republic of the Congo, Ghana, Kenya, Mozambique, Nigeria, Uganda, and Zambia) showing evidence of selective sweeps. H12 values reach maximum values (up to 1) in several cohorts. The model fits (Δi values ranging from 1,485 to 12,455) are among the highest observed across all loci in the *An. funestus* dataset, comparable to the strong signals observed in the *An. gambiae* complex.

#### Cyp9k1

The cytochrome P450 *Cyp9k1* was originally reported as being under selection in *An. gambiae* populations collected in Mali and Bioko island^60,61^. In the current study the *Cyp9k1* locus harbored selection signals in 37 cohorts distributed across Benin, Burkina Faso, Côte d’Ivoire, The Gambia, Ghana, Guinea, Mali and Uganda. Both *An. gambiae* and *An. coluzzii* populations display strong signals, with cohorts in Ghana, Burkina Faso, Mali, Cote D’Ivoire and Benin almost fixed for selective sweeps (H12 > 0.9). In *An. funestus, Cyp9k1* is commonly overexpressed and a mutation at codon 454 has been associated with pyrethroid resistance^14,62^. The locus shows strong signals of selection in populations from four countries (Democratic Republic of the Congo, Kenya, Nigeria, and Uganda), with H12 values ranging from 0.22 to 1.

#### Gste cluster

The glutathione S-transferase (GST) gene family plays a critical role in the detoxification of insecticides across many species and insecticide classes. In *An. gambiae*, increased expression and allelic variants of one member of this cluster, *GSTe2* (AGAP009194), has been implicated in DDT^19^ and pyrethroid^63^ resistance, as have orthologues in *Ae. aegypti* and *An. funestus*^20,64^. In vitro work has shown that this gene is involved in DDT metabolism^19^ and that a particular variant in this gene (114T) increases the efficiency of DDT metabolism irrespective of over-expression^19^. This locus exhibits signals of selection in 42 cohorts spanning 10 countries, in *An. gambiae* and *An. coluzzii*. H12 values are consistently high, peaking at 0.87. In *An. funestus*, an *Gste2*-L119F mutation enhances metabolism to DDT^20^ and is widespread^35^. The glutathione S-transferase cluster in *An. funestus* shows moderate to strong selection signals in populations from Benin, Democratic Republic of the Congo, Ghana, and Nigeria. H12 values range from 0.35 to 0.92, with the highest values observed in Benin.

#### Coeaexf

We also detected selection signals at the Coeaexf locus, recently implicated in resistance to the organophosphate, Pirimiphos-methyl^65^. *Coeae1f* and *Coeae2f* are orthologous to *Est3* and *Est2* in *Culex pipiens*, genes that were intensely studied in the late 20th century due to their role in organophosphate resistance. The Coeaexf locus is at 28.5Mbp on chromosome 2L, sitting within the region of the 2La chromosomal inversion. Other detoxification genes are in proximity to the two esterases, including the UGT AGAP006222. We detected signals in 13 cohorts from 7 countries (Benin, Burkina Faso, Côte d’Ivoire, Ghana, Guinea, Mali, and Tanzania), in *An. gambiae* and *An. arabiensis*. H12 values reach 0.6 indicating strong selection in some cohorts. No signals were observed at the orthologous locus in *An. funestus*.

### Selection signals at novel loci

#### KEAP1

At ~40.95 Mbp of chromosome 2RL, 400 kbp inside the 2Rd inversion, we detected signals in 28 cohorts, in *An. gambiae* and *An. coluzzii* from West and Central Africa, and from *An. arabiensis* in Malawi. H12 values are generally high in these cohorts peaking at 0.8 in Côte D’Ivoire. Our most promising candidate at this locus is *Keap1*, a member of the cnc-Maf-S pathway. This pathway is involved in the regulation of Cytochrome P450 and Glutathione S-Transferase families in *Drosophila* and other insects^66,67^. The pathway consists of three components, the *Maf-S* transcription factor, a cytoplasmic protein *cnc*, and an actin-binding ubiquitin ligase, *Keap1*. Using sequence homology, Ingham et al.,^68^ identified genes that putatively encode these proteins in *An. gambiae*. The *Keap1* gene (AGAP003645) is located at 40.92 Mbp on chromosome 2RL, close to the centre of the observed selection signals. Given the proven role of this protein in the regulation of metabolic resistance in insects, it is likely that this locus is responsible for the selection signal in this region. At the orthologous locus in *An. funestus,* we find a single selection signal in a cohort from the Ashanti region, Ghana.

#### RDGA / DGK

Another signal is present on the X chromosome, around 9.25 Mbp, in 11 cohorts of *An. gambiae* and *An. coluzzii* from West Africa. H12 values are mostly moderate except in *An. gambiae* from Côte D’Ivoire, where they peak at 0.94. The region at the centre of the selection signals in all three populations is relatively narrow (200 kbp), and centers over a single gene, AGAP000519. AGAP000519 is a diacylglycerol kinase, an enzyme responsible for the catalysis of diacylglycerol, predominantly involved in signalling. A potential mode of action is the regulation of synaptic transmission via the Gqα pathway, as shown in *C. elegans*^69,70^. Increased activity of diacylglycerol kinase could reduce synaptic excitability, potentially counteracting the neuronal hyperexcitation induced by insecticides. To our knowledge, there is no literature regarding the involvement of diacylglycerol kinases in insecticide resistance, and this may represent a previously unknown mediator of insecticide resistance.

This novel selection signal is also widespread in *An. funestus* populations, found in Cameroon, Democratic Republic of the Congo, Ghana, and Uganda. H12 values are moderate, ranging from 0.23 to 0.48. The presence of this signal in both major vector species provides additional evidence that this locus represents a previously unknown mechanism involved in insecticide resistance.

### Miscellaneous 34Mb signal

Extremely high H12, G123, and iHS values characterise a novel signal in East and central African *An. gambiae*, within the 2La inversion around 34.0 Mbp (Figure 2). H12 values are moderate, reaching 0.57 in *An gambiae* from Tanzania. This signal is present in 7 cohorts, originating from Uganda, Tanzania and Cameroon. The region putatively under selection is broad, encompassing approximately 1 Mbp, and was recently associated with resistance to deltamethrin nets in an amplicon sequencing study^33^. The sweep is present on the inverted form^71^, and so its spread may be limited to the environmental niche of each karyotype.

**Figure 2.**
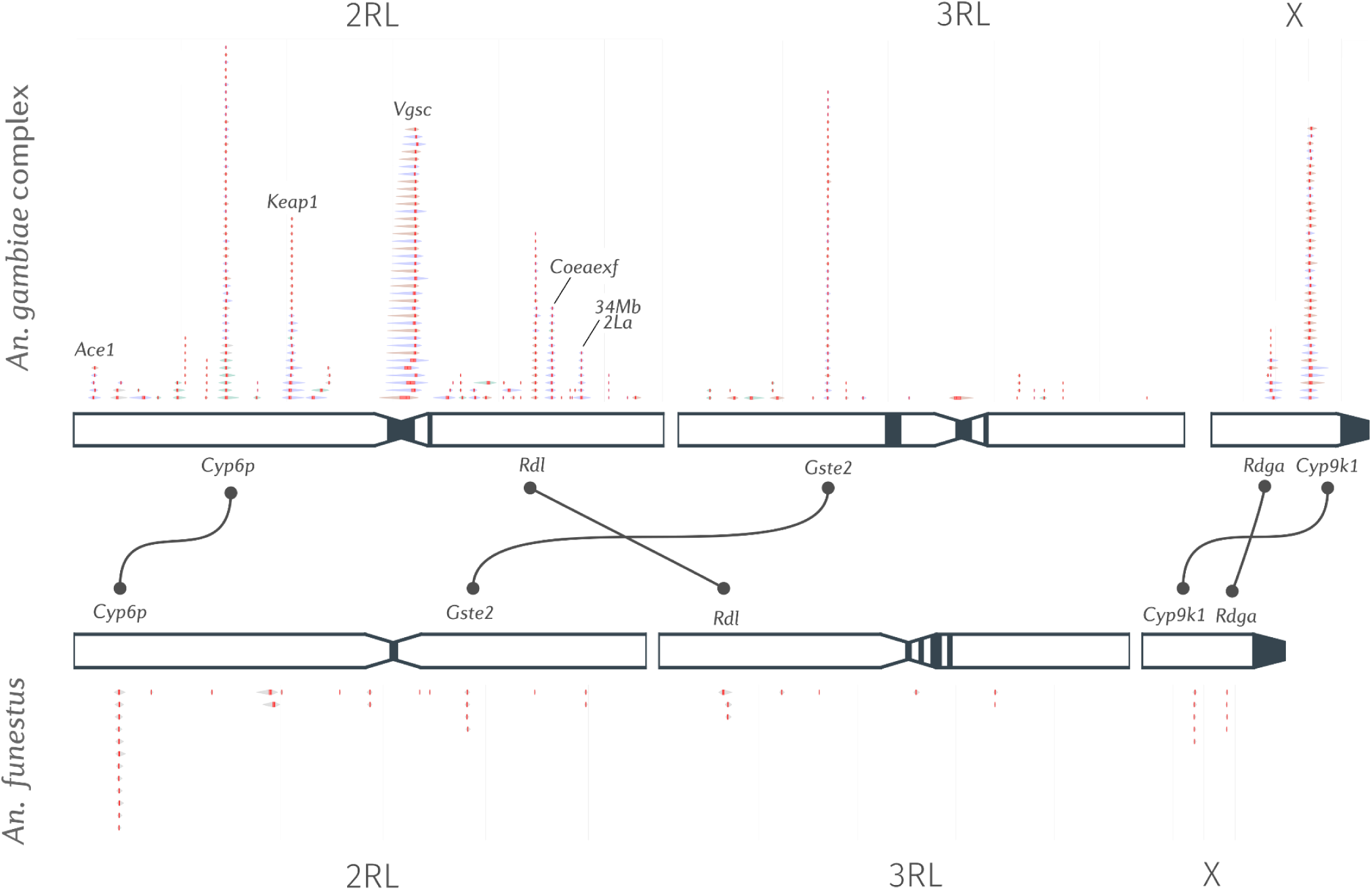
Summary of H12 selection signals across the *An. gambiae complex* and *An. funestus* in sub-Saharan Africa. H12 selection signals from different cohorts are stacked on top of each other, highlighting both known resistance loci and novel selection signals. Each diamond shape is a H12 signal in a single cohort called by the signal detection algorithm. The width of a signal indicates its breadth.

At the peak of the signals is AGAP006541, encodes a protein containing both a DNA-binding domain and an aminotransferase domain, which could suggest a role in transcriptional regulation of metabolic pathways. However, owing to the broad nature of the signal and the large number of genes present in the swept locus, a clear driver is not evident in this case. We also examined expression data from a recent transcriptomic meta-analysis^14^, however, no genes in this genomic region showed systematic bias towards over-expression in resistant strains of *Anopheles gambiae sl* (Supplementary Figure 1).

## Discussion

In this manuscript we report the results from genome-wide scans for positive selection in 4,306 mosquitoes sampled across twenty-one African countries. We have created an open-source, accessible web platform, the Malaria Vector Selection Atlas, that systematically highlights genomic regions that may be threatening chemical-based vector control. The selection atlas integrates multiple selection statistics (H12, G123, and iHS) with a novel peak-finding algorithm to identify driving loci. Importantly, it is implemented as an automated, reproducible computational workflow, which enables continuous updates as new whole-genome sequence data become available. The web interface is accessible for researchers and vector control stakeholders without specialized bioinformatic expertise. By presenting complex genomic data through intuitive visualizations, we hope the resource can help to bridge the gap between genomics and practical vector control operations.

Within the selection atlas, we present selection alerts - a curated list of loci putatively threatening vector control, containing both known and novel regions. As future datasets reveal additional loci of concern, we will continue to expand this resource while advancing amplicon-based approaches as a cost-effective strategy for routine insecticide resistance monitoring^33^. Our ultimate goal is to identify and characterise selective sweeps at every locus under selection, creating a comprehensive database of haplotypes, their defining variants, and an accompanying nomenclature. This database can then feed into existing targeted surveillance platforms, such as Ag-vampIR^33^, and implemented with optimised sampling frameworks^72^ to enable more complete and effective predictive monitoring of the resistance phenotype.

Currently, the *An. gambiae s. l.* vector sequencing data is all mapped to an outdated reference genome (PEST) and undoubtedly, especially for *An. arabiensis*, this results in regions of the genome that have mapping quality issues. Future revision of the selection atlas may split *An. gambiae*, *An. coluzzii*, and *An. arabiensis* as these genetic variation datasets get mapped to their appropriate reference genomes, which now exist in the public domain.

The recent introduction of dual-active ingredient LLINs and novel IRS insecticides is introducing new selection pressures that will shape mosquito adaptation in coming years. By updating the resource as new data become available, we can track the evolutionary response of these vectors in near real-time. By providing an open-source, updateable resource, we hope the selection atlas can support evidence-based decision making and proactive resistance management. As we enter a new era of novel chemistries, we hope this resource will play a role in preserving the effectiveness of vector control tools and sustaining progress toward malaria elimination.

## Materials and Methods

### Data collection and sample processing

We analyzed whole-genome sequence data from 4,306 mosquitoes collected across twenty-one African countries as part of phase 3 of the *Anopheles gambiae* 1000 Genomes Project, the *Anopheles funestus* 1000 genomes project, and the Vector Observatory^34,35^. Individual mosquitoes were collected as adults or larvae between 1994 and 2018. DNA extraction, sequencing, variant calling, and phasing were performed using standardized protocols following previously established methods^73^. In brief, high-coverage (30×) whole-genome sequencing was performed, reads mapped with BWA-MEM, genotypes called with GATK and haplotypes phased with SHAPEIT2.

### Cohort assignment and filtering

Individual mosquitoes were grouped into cohorts based on the geographic district (administrative level 2) and the quarter of the year in which they were collected. To ensure statistical robustness for selection analyses, we only included cohorts with at least 15 individuals. For cohorts with more than 200 mosquitoes, we randomly downsampled to 200 individuals to standardize computational approaches and minimize biases from uneven sample sizes.

### Genome-wide selection scans

For each cohort, we computed three complementary genome-wide selection statistics: H12, G123, and integrated haplotype score (iHS). These methods detect different signatures of positive selection, with iHS identifying incomplete sweeps, H12 capturing both hard and soft sweeps, and G123 providing reliability in genomic contexts where phasing is challenging. Together, they offer a reliable framework for detecting and validating regions under positive selection.

H12 is a haplotype homozygosity-based statistic that provides a measure of haplotype diversity within a population^36^. A high H12 value indicates unusually low haplotype diversity, expected in genomic windows where recent selective sweeps containing beneficial alleles have spread to moderate or high frequency. For each cohort, we first performed a calibration step to determine the appropriate window size for H12 analysis. We tested window sizes of 200, 500, 1000, 2000, 5000, and 10000 bp on chromosome arm 3L, which serves as a control region with relatively few known resistance loci. We selected the smallest window size for which the 95th percentile of H12 values was below 0.1, ensuring that background variation would not produce excessive false positive signals. Using the calibrated window size for each cohort, we calculated H12 values in sliding windows.

G123 is analogous to H12 but applied to unphased diplotypes rather than haplotypes^37^. This approach captures similar signatures of selection without requiring haplotype phasing, making it particularly useful where phasing may be challenging, such as at the sites of copy number variation. As with H12, we performed window size calibration separately for each cohort before computing genome-wide G123 values.

The integrated haplotype score (iHS) was computed for each cohort using all segregating SNPs above a minor allele frequency of 5%^74^. A high absolute value of iHS at a given SNP indicates that one allele has experienced recent positive selection because it is found on longer haplotypes than the alternative allele. Although iHS was originally defined as a comparison between derived and ancestral alleles, to avoid bias due to errors in allele polarization, we compared alternate with reference alleles and then took the absolute value. We used a fixed window size of 100 SNPs for the iHS analysis.

### Signal detection algorithm

To systematically identify genomic regions under selection, we developed a peak-finding algorithm that accounts for the extended linkage disequilibrium surrounding loci under selection. Our approach is based on fitting exponential decay curves to the selection scan data using non-linear least squares regression, which captures the characteristic decay pattern of selection signals. For each cohort and chromosome, we applied the following procedure to the H12 data: We first pre-processed the H12 values using a Hampel filter to remove outliers that might represent technical artifacts. We then converted physical positions (bp) to genetic positions (cM) using a genetic map of An. gambiae derived from previously published recombination maps. Next, we scanned the genome in 1 cM intervals, fitting three models at each location: a skewed exponential decay model with parameters for center, amplitude, decay rate, skew, and baseline; a Gaussian peak model; and a null constant model representing no selection. For each window, we computed the difference in Akaike Information Criterion (ΔAIC) between the best-fitting alternative model and the null model. Windows with ΔAIC > 1000 and peak H12 value > 0.1 were recorded as candidate selection signals. Finally, we removed redundant overlapping signals, retaining only the one with the highest ΔAIC.

### Selection atlas web resource

To make the results accessible to the broader research and public health communities, we developed an open-source web resource for exploring the selection scans. The resource was implemented as an automated computational workflow using the workflow manager Snakemake^75^. The workflow is split up into two components: A genome-wide selection scan (GWSS) workflow that processes the raw data and performs the statistical analyses, and a site-building workflow that compiles the results into an interactive web resource. Results from a given run are stored in Google Cloud, to allow contributors to download the data and build the site without re-running the entire GWSS workflow.

The selection atlas web interface was built using Jupyter Book^76^ and includes interactive visualizations of selection scans, geographic mapping of cohorts, and information about selection signals at known and novel resistance loci. The interface allows users to explore results by chromosome and cohort. We classified selection signals into “alerts” based on their replication across at least 5 cohorts.

## Data availability

All code and configuration files for the analysis workflow and web resource are available in the public GitHub repository (github.com/anopheles-genomic-surveillance/selection-atlas). The web resource is available at anopheles-genomic-surveillance.github.io/selection-atlas/ and whole genome sequence data for all cohorts is available through the malariagen_data python API.

## Acknowledgments

This work was supported by the National Institute of Allergy and Infectious Diseases (NIAID R01-AI116811 to M.J.D) and the Medical Research Council (MR/T001070/1 to M.J.D., MR/P02520X/1 to M.J.D). The latter grant is a UK-funded award and is part of the EDCTP2 program supported by the European Union. M.J.D. was supported by a Royal Society Wolfson Fellowship (RSWF\FT\180003). This study was supported by the MalariaGEN Vector Observatory which is an international collaboration working to build capacity for malaria vector genomic research and surveillance, and involves contributions by the following institutions and teams. Wellcome Sanger Institute: Lee Hart, Kelly Bennett, Anastasia Hernandez-Koutoucheva, Jon Brenas, Menelaos Ioannidis, Chris Clarkson, Alistair Miles, Julia Jeans, Paballo Chauke, Victoria Simpson, Eleanor Drury, Osama Mayet, Sónia Gonçalves, Katherine Figueroa, Tom Madison, Kevin Howe, Mara Lawniczak; Liverpool School of Tropical Medicine: Eric Lucas, Sanjay Nagi, Martin Donnelly; Broad Institute of Harvard and MIT: Jessica Way, George Grant; Pan-African Mosquito Control Association: Jane Mwangi, Edward Lukyamuzi, Sonia Barasa, Ibra Lujumba, Elijah Juma. The authors would like to thank the staff of the Wellcome Sanger Genomic Surveillance Unit and the Wellcome Sanger Institute Sample Logistics, Sequencing and Informatics facilities for their contributions. The MalariaGEN Vector Observatory was supported by funding awarded to Dominic Kwiatkowski and Mara Lawniczak from Wellcome (220540/Z/20/A, “Wellcome Sanger Institute Quinquennial Review 2021-2026′) and funding awarded to Dominic Kwiatkowski from the Bill and Melinda Gates Foundation (INV-001927). The Pan-African Mosquito Control Association’s participation was funded by the Bill and Melinda Gates Foundation (INV-031595).

---

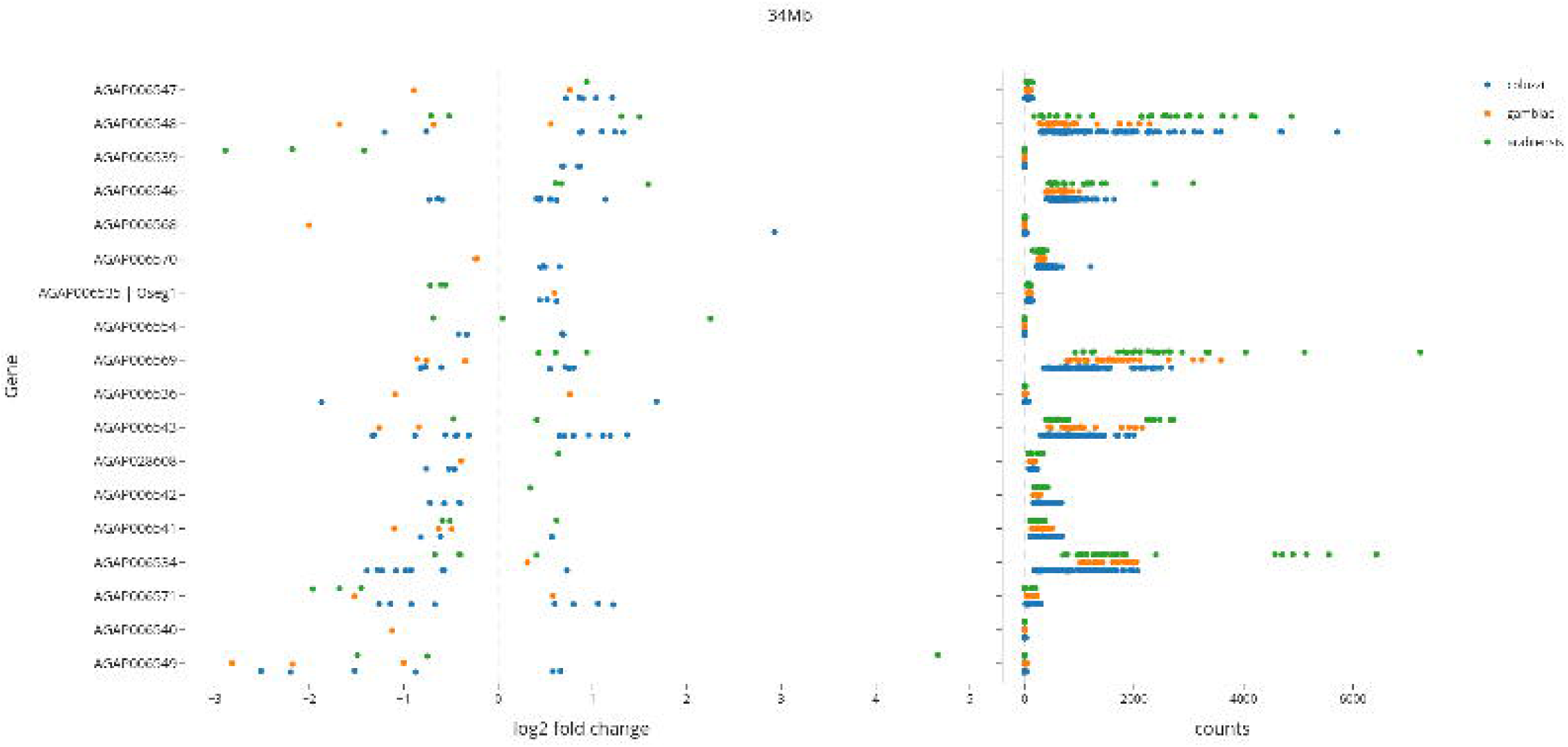

